# Whole-exome sequencing identified rare variants associated with body length and girth in cattle

**DOI:** 10.1101/287474

**Authors:** Yan Chen, Min Du, Yining Liu, Xue Gao, Wengang Zhang, Ling Xu, Lupei Zhang, Huijiang Gao, Lingyang Xu, Bo Zhu, Min Zhao, Junya Li

**Author notes:** Corresponding authors. Junya Li, Institute of Animal Science (IAS), Chinese Academy of Agricultural Sciences (CAAS), No. 2 Yuanmingyuan West Road, Beijing, 100193, China.; Min Zhao, Faculty of Science, Health, Education and Engineering, University of the Sunshine Coast, Bldg I, Rm 1.11B, ML 12.; Yan Chen, Institute of Animal Science (IAS), Chinese Academy of Agricultural Sciences (CAAS), No. 2 Yuanmingyuan West Road, Beijing, 100193, China.

## Abstract

Body measurements can be used in determining body size to monitor the cattle growth and examine the response to selection. Despite efforts putting into the identification of common genetic variants, the mechanism understanding of the rare variation in complex traits about body size and growth remains limited. Here, we firstly performed GWAS study for body measurement traits in Simmental cattle, however there were no SNPs exceeding significant level associated with body measurements. To further investigate the mechanism of growth traits in beef cattle, we conducted whole exome analysis of 20 cattle with phenotypic differences on body girth and length, representing the first systematic exploration of rare variants on body measurements in cattle. By carrying out a three-phase process of the variant calling and filtering, a sum of 1158, 1151, 1267, and 1303 rare variants were identified in four phenotypic groups of two growth traits, higher/ lower body girth (BG_H and BG_L) and higher/lower body length (BL_H and BL_L) respectively. The subsequent functional enrichment analysis revealed that these rare variants distributed in 886 genes associated with collagen formation and organelle organization, indicating the importance of collagen formation and organelle organization for body size growth in cattle. The integrative network construction distinguished 62 and 66 genes with different co-expression patterns associated with higher and lower phenotypic groups of body measurements respectively, and the two sub-networks were distinct. Gene ontology and pathway annotation further showed that all shared genes in phenotypic differences participate in many biological processes related to the growth and development of the organism. Together, these findings provide a deep insight into rare genetic variants of growth traits in cattle and this will have a promising application in animal breeding.

## 1. Introduction

Beef cattle production plays an important role in Chinese agribusiness. According to the statistics by China’s Ministry of Agriculture, beef production has five years of continuous growth, up to 83% increase by 2015 than 2011. Many breeds specialized in meat production are reared in China, but Simmental breed accounts for more than 70% of beef-producing herds. Growth traits are traditionally included in selection criteria in beef cattle breeding programs. Body linear measurements, specifically body length and girth, have been shown to be useful predictors of cattle liveweight (Enevoldsen and Kristensen, 1997; Lukuyu et al., 2016).

With the genome sequencing of common cattle, more and more interaction between genes and environments are determined to steadily increase the quality and quantity of diary and cattle meat in global (Zimin et al., 2009). Recently, a few novel genetic loci associated with quantitative traits have been detected by genome-wide association studies in bovine and other livestock species using high or low-density SNP array and sequencing approach. For instance, eight significantly SNPs, seven potential genes, and two most important quantitative trait loci regions were identified for improved lactation persistency in Holstein cattle (Pertille et al., 2017). In addition, by focusing on beef cattle, numerous SNPs, genes and haplotype blocks were discovered associated with growth (Jahuey-Martinez et al., 2016; Sorbolini et al., 2017). In our previous study, pathway-based GWAS method was applied to identify novel loci and candidate genes of complex quantitative economically important traits in beef cattle (Fan et al., 2015; Xia et al., 2017). We further identified the DCAF16-NCAPG region as a susceptibility locus for average daily gain in Chinese Simmental cattle (Zhang et al., 2016b).

In general, the current association studies have been performed for identification of common variants to explain the complex traits (Bush and Moore, 2012; Hirschhorn and Daly, 2005). Despite their value, the GWAS-based studies may have some problems like population stratification, reproducibility and high false positive. Furthermore, the functional validation of the GWAS result is the major challenge. In addition, the commercial designed SNP arrays were mainly focused on common variants with a high frequency in the population. Thus, the studies failed to discover those low-frequency/rare genetic variants that may affect functional properties, especially in certain specific breeds (Clayton et al., 2005; Donnelly, 2008). To overcome these shortcomings, there is a trend to integrate more rare variations for follow-up functional validation. For example, whole-genome sequencing analysis has demonstrated the advantage to map rare genetic variants that affect quantitative traits in cattle and other livestock with complex familial relationships (Zhang et al., 2016a). In addition, rare causative variants could improve genomic prediction, but careful selection of markers was needed (van den Berg et al., 2016).

It is worth noting that the majority studies focused on dairy cattle instead of beef cattle. The comprehensive evaluation of rare variants in beef cattle is still lacking. By using the whole exome sequencing (WES) to economically important traits, we performed the first study to explore the rare variants associated with growth traits in Simmental cattle. The results provide a better genomics prediction and more accuracy of gene marker selection for complex traits in beef cattle.

## 2. Materials and methods

### 2.1 Cattle population, phenotypes, genotyping and GWAS for body measurements in Simmental cattle

Since 2008, we established the Simmental cattle population in Ulgai, Inner Mongolia, China. A total of 1141 Simmental beef cattle born between 2008 and 2014. Phenotypic data including growth traits like body length, height, hip height, heart girth, and abdominal girth was collected at regular intervals. For each animal, 10 mL of venous blood was collected from the jugular vein and then stored at –20°C. The DNAs were extracted from the blood samples and were genotyped using Illumina BovineHD BeadChip. Genome wide association studies (GWASs) analyses were implemented based on mixed linear model as we described previously (Xia et al., 2016; Zhang et al., 2016c).

### 2.2 Sample collection and whole exome sequencing

To investigate a systematic survey of growth traits in beef cattle, body length and body girth were chosen as two parameters for stature in this study. The two traits in accordance with the phenotype value were divided into high and low groups, respectively. The criteria were: i) 18-month-old phenotypic data were used at which time a calf almost reached maturity. ii) each of high or low group had five samples, randomly selecting from the top 20 or bottom 20 ranked-list of phenotype data.

Approximately 1μg to 2μg of DNA was obtained and quantified using Blood DNA Kit, Nanodrop and Qubit fluorometer. Exome capture was accomplished using the Agilent SureSelect All Exon Bovine (54Mb design based on University of Maryland build 3.1, covering coding regions as well as miRNA and SNP targets). Sequencing was performed using the Illumina HiSeq4000 system at an average coverage depth of over 150X (Shanghai OE Biotechnology Co., Lt, China).

### 2.3 The bioinformatics pipeline for variant calling and filtering

The second and third phases for the WES analysis were to detect, annotate and filter genetic variants for each cattle sample. To this aim, we filter out those low-quality reads by using NGSQC-Toolkit (v2.3.3). Briefly, we firstly removed those raw reads with Q20 value less than 70%, which represented the ratio of bases with probability of containing no more than one error in 100 bases. Then we trimmed reads shorter than 70 bases afterwards to obtain high-quality reads. Next, all high-quality paired reads were extracted with SAM flags in the raw BAM files using Samtools (version 0.1.19) (Li et al., 2009). In each sample, the short reads were aligned to the bovine reference genome, University of Maryland build (UMD 3.1) using BWA-mem (version 0.7.5a) (Jiang et al., 2010). The mapped reads were sorted and indexed by using Samtools (version 0.1.19) (Li et al., 2009). The mpileup command in Samtools toolkit (version 0.1.19) was applied to call single nucleotide variant (SNV), and insertion/deletion (INDEL) variants using the BAQ (Base Alignment Quality) with cut-off of 13 and minimum/maximum read depth cut-off of 1/8000.

### 2.4 The functional enrichment and QTL analysis

Through capture sequencing, hundreds of rare nonsynonymous variants were identified to be associated with four trait groups, and we hypothesized that there might be commonalities or similarities in the biological functions of the affected genes for body growth traits. To explore those potential central pathways involving the cattle growth, we run gene ontology (GO) and KEGG pathway enrichment analysis by inputting those mutated genes against all the cattle protein-coding genes as background. Comparing our 886 mutated genes, the statistically overrepresented GO terms and KEGG pathways were listed using BovineMine (Elsik et al., 2016). The biological processes and interrelated genes were graphically displayed as we described previously (Zhao et al., 2009; Zhao et al., 2013). In addition, the cattle QTL information was downloaded from animal QTL database (Hu et al., 2013) and mapped to genes according to the official gene symbols.

### 2.5 Sub-network extraction for the genes with genetic changes from a co-expression network in cattle

To elucidate genetic variations on a global scale, we mapped the variants to the corresponding genes and constructed the gene co-expression network. The gene co-expression network was built based on gene profiles derived from 92 different tissues from Bovine Genome Database (BGD) (Elsik et al., 2016). These data are generated by single-end RNAseq with 100 bp reads running on Illumina HiSeq 2000. By mapping to genome, BGD has normalized the read counts for each tissue. The normalized FPKM (Fragments Per Kilobase of transcript per Million mapped reads) value was used to calculate the co-expression based on WGCNA (Weighted Gene Correlation Network Analysis) methods. In total, we identified 72,306 pairs of genes that are co-expressed over 92 tissues (Chen et al., 2017). To explore the biological mechanisms related to genes that may exert the consistent effects in high phenotype groups for body girth and length traits (BG_H and BL_H), we extracted the protein-protein interactions between 62 identified genes with the remaining cattle genes. The similar network mapping approach was applied to the 66 genes shared between low phenotype groups (BG_L and BL_L groups). For co-expression based interactions, we used the Steiner minimal tree algorithm implemented in our previous studies to extract a sub-network related to the input genes (Kong et al., 2013). In this algorithm, all inputted genes were mapped to the co-expression based interactome. Finally, a minimum sub-network with inputted genes connected by shortest path was produced. The final network visualization was performed using Cytoscape (Shannon et al., 2003).

## 3. Results

### 3.1 Genome-wide association studies (GWAS) for body measurements in Simmental cattle

Given the economic importance of growth traits in beef cattle, genome-wide association studies (GWASs) for body measurements including five traits (body length, height, hip height, heart girth, and abdominal girth) were performed in 1141 Simmental cattle using Illumina BovineHD BeadChip. However, we found there were no SNPs exceeding significant level (*P*< 5×10^−7^) for the five traits related to body measurements. Therefore, these results indicate common variants of small effect in polygenic model may not be tagged by genotyping arrays. Manhattan plots and quantile-quantile (Q–Q) plots of genome-wide association results were showed in Figure 1.

**Figure 1.**
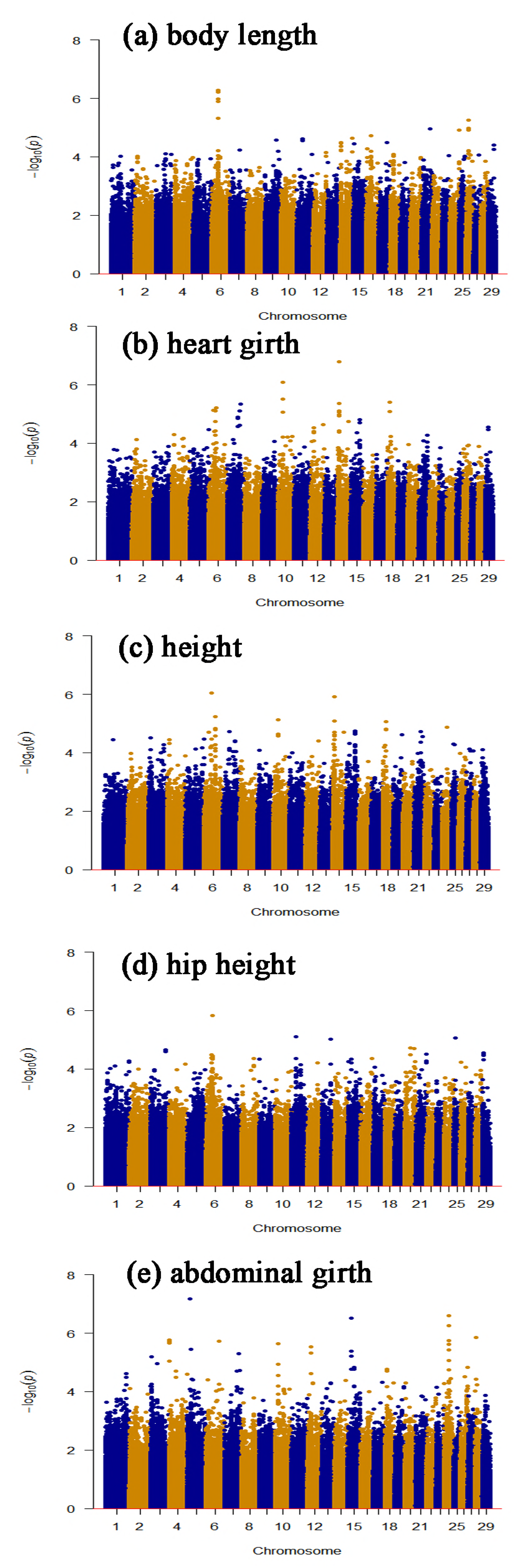
Manhattan plots and quantile-quantile (Q–Q) plots of genome-wide association results for body measurement in Simmental cattle. (a) body length, (b) heart girth, (c) height, (d) hip height, (e) abdominal girth.

### 3.2 The sample information and an overview for whole exome sequencing classified into four groups related to body length and girth

Since we did not find any significant SNPs and enriched gene regions associated with Simmental body measure traits based on the GWAS method, we further sought to identify rare and novel genetic variants with large effects on body size measurement to predict cattle liveweight and visual assessment to meat production, growth and development, and nutritional. Taking into account the contributions of body length and girth, the 20 samples we collected were classified into four distinct trait groups: high body girth (BG_H), low body girth (BG_L), high body length (BL_H), and low body length (BL_L), based on sorting the value of phenotypic data. The detail characteristics were summarized (Table S1). To identify the potential rare variants associated with body girth and length, we conducted a three-phase analysis process for whole exome sequencing and follow-up bioinformatics analysis (Figure 2). By using the Agilent SureSelect All Exon Bovine toolkit (54 Mb design covering protein-coding regions and SNP targets), all the samples were loaded to Illumina HiSeq4000 for sequencing. In the second phase, we conducted the bioinformatics data processing step by step (see methods for detail parameters): i) data quality control to remove low quality reads; ii) map to Bovine UMD 3.1 genome; iii) call and annotate the genetic variants. In our last analysis stage, we started from all the identified genetic variants in the 20 samples. After excluding those synonymous and common variants in the population, we focused on the rare non-synonymous variants shared in different trait groups. The set-based analyses were conducted to identify key rare variants shared in different trait combination.

**Figure 2.**
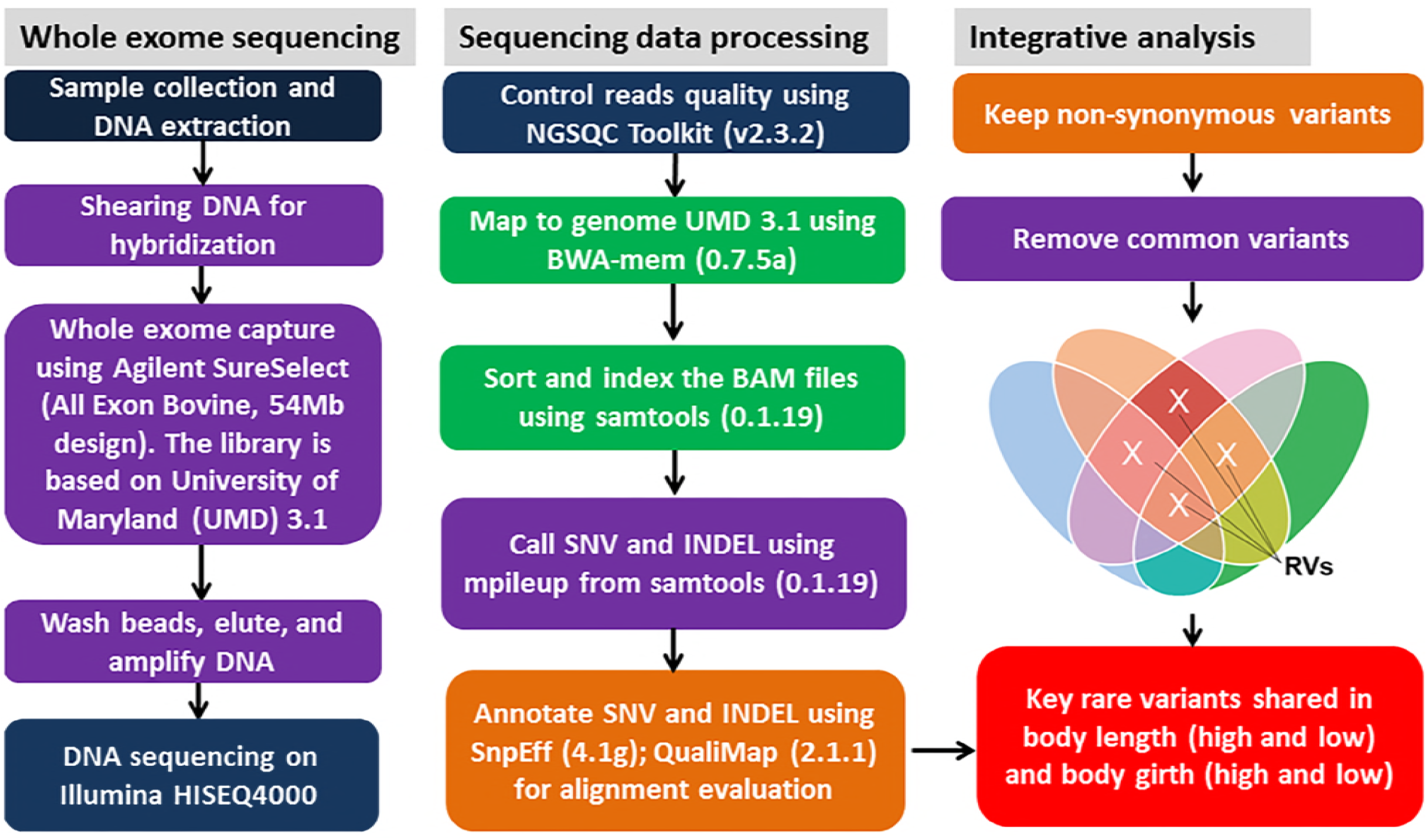
Whole-exome sequencing (WES) analysis pipeline. The WES analysis occurred through three phases. In phase 1, all the exome DNA in the collected samples were extracted, sheared and enriched by Agilent SureSelect for sequencing on Illumina HiSeq4000. In phase 2, all the raw reads underwent quality controlling, genome mapping, variant calling and annotation. In phase 3, only those highly likely to affect protein quantity or function (nonsynonymous exonic single-nucleotide variants [i.e., missense or nonsense], insertions and deletions, splice variants) were selected for further analysis. Among these non-synonymous variants, those common variants mapped to dbSNP were excluded to yield candidate rare variants. Following exclusionary quality-control filter, only rare non-synonymous variants were compared in all four distinct groups: higher body girth (BG_H), lower body girth (BG_L), higher body length (BL_H), and lower body length (BL_L). INDEL = insertion and deletion; SNV = single-nucleotide variants; RV =rare variant.

### 3.3 The variant filtering and functional analysis for the nonsynonymous rare variants

A step-by-step variant filtering approach was applied to narrow down the candidate genetic variants associated with four trait groups (BG_H, BG_L, BL_H and BL_L). For example, we identified 45,896 variants shared by the five samples from BG_H group (Figure 3A, Table S2). By filtering out the synonymous variants, we found a total of 22,788 nonsynonymous variants. By further removing the common variants with SNP IDs from the dbSNP database, we narrowed down the list to 1158 rare variants, which could be mapped to 299 genes. To obtain precise genetic variants for different traits, the same variants filtering was applied to those genetic variants shared by all the samples in the other three trait groups. Finally, we identified 302, 263, and 283 genes associated with the rare variants for BG_L, BL_H, and BL_L groups respectively.

**Figure 3.**
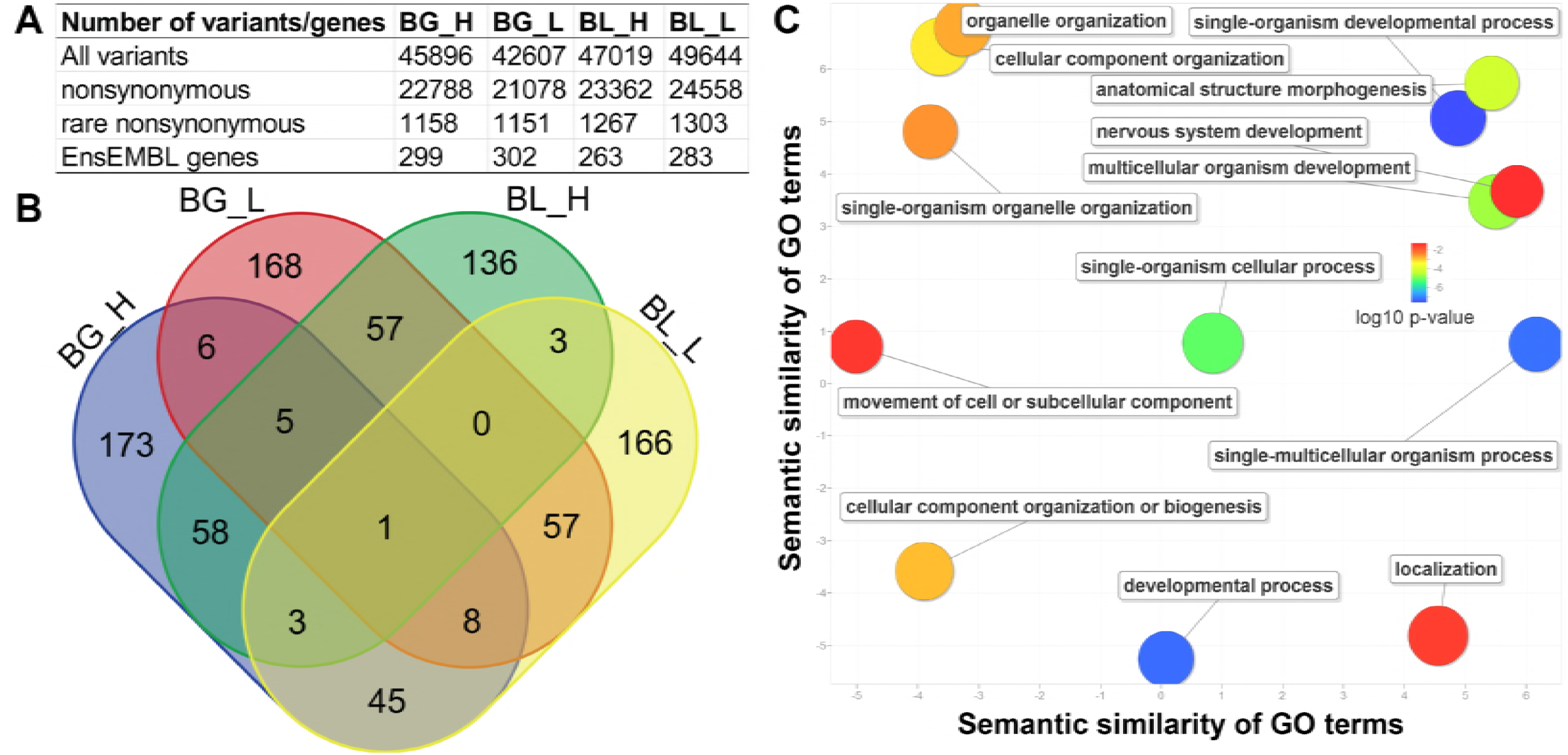
The variant filtering and functional analysis of genes associated with four different groups. (A) The variant filtering statistics for four distinct groups: higher body girth (BG_H), lower body girth (BG_L), higher body length (BL_H), and lower body length (BL_L). (B) The relationship for all the 886 Ensemble genes associated with four different traits was presented. (C) Gene ontology (GO) enrichment analysis. The scatterplot showed the gene ontology (GO) cluster representatives for all the 886 genes in a two-dimensional space derived by applying multidimensional scaling to a matrix of the GO terms’ semantic similarities. Bubble colour indicated the corrected *P*-values (bubbles of more significant terms were blue).

As shown in Figure 3B, the four trait groups shared some associated genes. Basically, there were relative less shared genes between two BG groups (BG_H and BG_L) and BL groups (BL_H and BL_L). To provide an overview for those genes, we performed functional enrichment on all the identified 886 unique genes (union analysis, Table S3). The interesting finding was that these genes were more likely to be involved in various developmental processes (Figure 3C, Table S4), including single-organism development process (adjusted *P*-value = 2.71E-08), developmental process (adjusted *P*-value = 5.72E-08), anatomical structure morphogenesis (adjusted *P*-value = 2.40E-06), multicellular organism development (adjusted *P*-value = 7.23E-06), and nervous system development (adjusted *P*-value = 4.86E-02). In addition, we also identified a number of organelle organization related functional GO terms, including cellular component organization (adjusted *P*-value = 5.79E-04), cellular component organization or biogenesis (adjusted P-value = 1.37E-03), organelle organization (adjusted *P*-value = 1.94E-03), single-organism organelle organization (adjusted *P*-value = 3.00E-03), and movement of cell or subcellular component (adjusted *P*-value = 3.67E-02). The further KEGG pathway analysis revealed that these genes were significantly over-represented in six pathways, including Laminin interactions (adjusted *P*-value = 1.13E-05), Collagen formation (adjusted *P*-value = 3.12E-04), Extracellular matrix organization (adjusted *P*-value = 1.92E-03), Collagen biosynthesis and modifying enzymes (adjusted *P*-value = 1.50E-02), Assembly of collagen fibrils and other multimeric structures (adjusted *P*-value = 4.16E-02), and Rho GTPase cycle (adjusted *P*-value = 4.61E-02).

### 3.4 Genetic and network difference between high and low body measurement phenotype groups in cattle

Our focus on those samples with the same characteristics of a trait can help identify the shared genetic variations. For instance, we identified 45,896 and 42,607 shared variants among five samples from high and low phenotype groups of body girth trait (BG_H and BG_L), respectively. By intersecting these two set, we characterized 17,466 and 14,177 genetic variations unique for BG_H and BG_L samples (Figure 4). The same intersecting analysis was applied to two groups of body length trait (BL_H and BL_L), and 14,774 and 17,399 genetic variations were found unique to BL_H and BL_L respectively.

**Figure 4.**
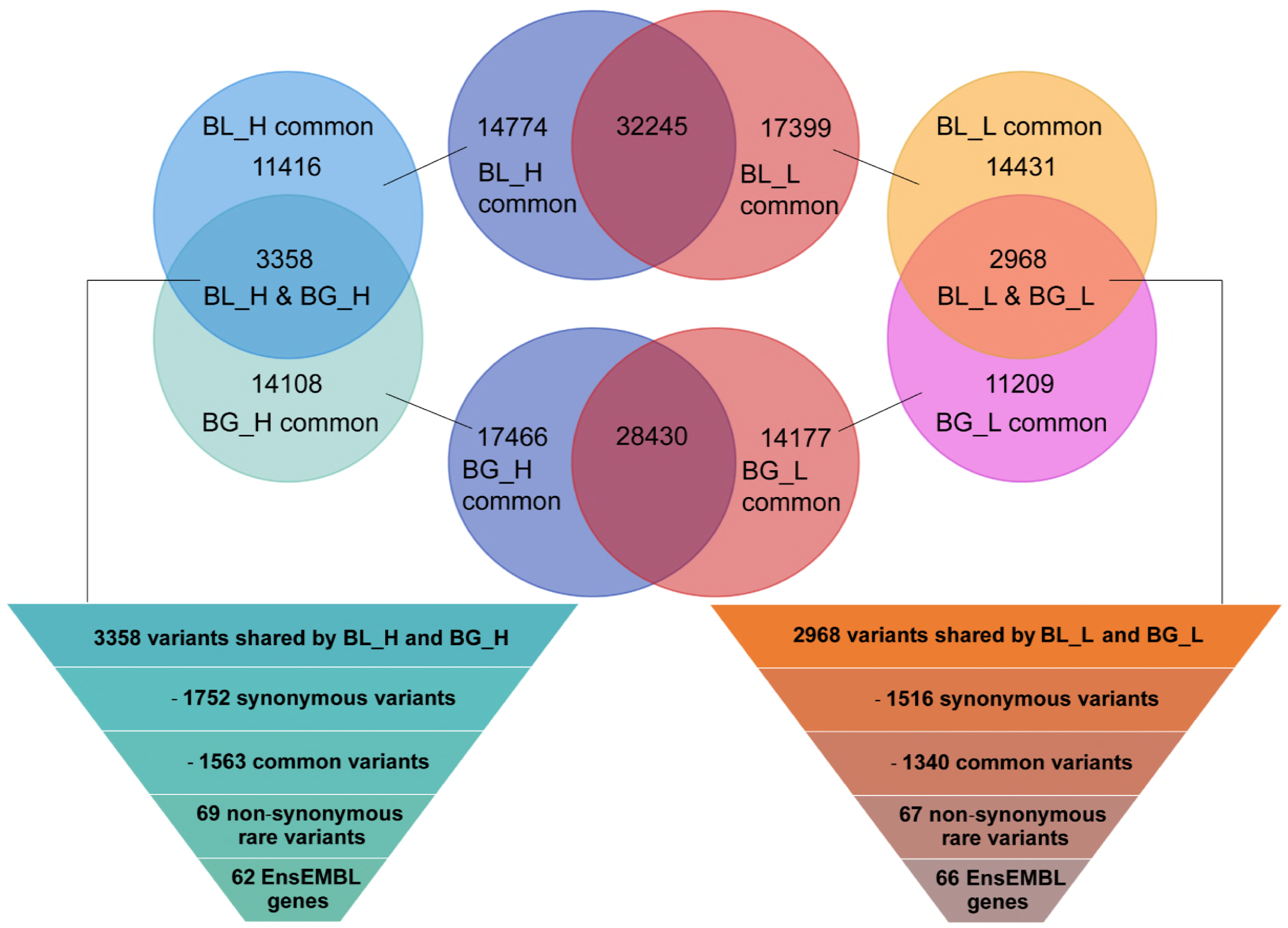
The step-by-step variant filtering for identification of rare genes associated with higher and lower phenotype in body girth and body length. To identify shared genetic variants in higher body girth (BG_H) and higher body length (BL_H), we started with 14,774 BL_H and 17,466 BG_H unique variants. The 3358 shared variants were further filtered and mapped to 62 Ensemble genes. The same filtering pipeline was applied to lower body girth (BG_L) and lower body length (BL_L) and 66 shared genes were obtained.

Regardless of the cause, traits with high degree of genetic correlation (whether positive or negative) are generally considered to be under the control of the genes with linkage or pleiotropy. Since BG and BL are highly positive genetic related traits, we further focused on those genetic variations shared by the similar phenotype with higher value in BG_H and BL_H samples, which may be associated with growth and gain of weight of cattle. As shown in Figure 4, there were a total of 3358 genetic variants shared between BG_H and BL_H groups. The same intersecting analysis identified 2968 variants shared by the lower phenotype groups in BG_L and BL_L samples. By conducting step-by-step variant filtering, we first harvested 69 nonsynonymous rare variants for both high groups (BG_H and BL_H), which could be mapped to 62 genes (Table S5_a). For example, *BIN2* and *ERAP1* had rare mutations at donor or acceptor sites in the splice region. There were two rare mutations were located in *ANGEL2* and *SPATA22* occurred in the 5’ UTR; and two insertion/deletion in the corresponding genes, *CLSPN* and *MUC3A*, respectively caused the disruption of the open reading frame. Most importantly, we identified four missense variants in the coding region of four genes, *SCN5A*, *BOLA-DRB3*, *FADS2* and *ENSBTAG00000046327*, which may lead to the functional changes or even deficiencies of the encoding proteins. The same approach was applied to BG_L and BL_L groups and identified 66 genes with 67 nonsynonymous rare variants shared between these two low groups (Table S5_b). For example, *EDN3* and *AKR1E2* had rare mutations at donor or acceptor sites in the splice region; seven rare mutations induced to frameshift of the coding region in *IGFN1*, *LPIN1* and other five genes. It is noted that one missence mutation occurred in a novel gene, ENSBTAG00000007696, which encode an uncharacterized protein involving endoplasmic reticulum (ER) to Golgi vesicle-mediated transport and protein localization to pre-autophagosomal structure. Gene ontology and pathway annotation analysis showed that all shared 62 and 66 genes in phenotypic differences participate in many biological processes related to the growth and development of the organism, such as regulation of cell differentiation and proliferation (*EDN3*), growth factor activity (*VEGFC*), fatty acid biosynthetic and metabolic process (*FADS2*), bone morphogenesis (*GLG1*), and signal transduction (*MRAS*, *TYK2* and *GSK3B*).

The co-expressed genes in cell may mediate similar biological function and form connected functional modules to play a pivotal role. Previous studies revealed that the whole co-expression network in cattle (‘interactome’) (1) follows a power-law degree distribution, (2) exhibits the small world behaviour and (3) tends to be modular (Beiki et al., 2016; Chen et al., 2017; Ghorbani et al., 2015). Therefore, identification of sub-networks with special characteristics using graphical approaches can also lead to biologically relevant insights. In general, the densely-interconnected gene-gene pairs in a global co-expression network often correspond to functionally related groups of genes that can be defined as modules. To further explore the biological function and improve a systems-level understanding of the relations for the 62 (shared by two high groups in BL_H and BG_H) and 66 genes (shared by two low groups in BL_L and BG_L) from strict filtering, we performed the gene co-expression based network analysis on the two gene lists separately. By using a module extraction algorithm, we connected those input genes as more as possible. Therefore, there were two types of genes in the final output network: the genes from our interesting genes and the linker genes to connect those genes. As shown in Figure 5, the reconstructed sub-network specific for the 62 genes shared by the higher phenotype groups (BL_H and BG_H) was distinct from the 66 genes shared by the lower phenotype groups (BL_L and BG_L) in terms of their gene co-expressions.

**Figure 5.**
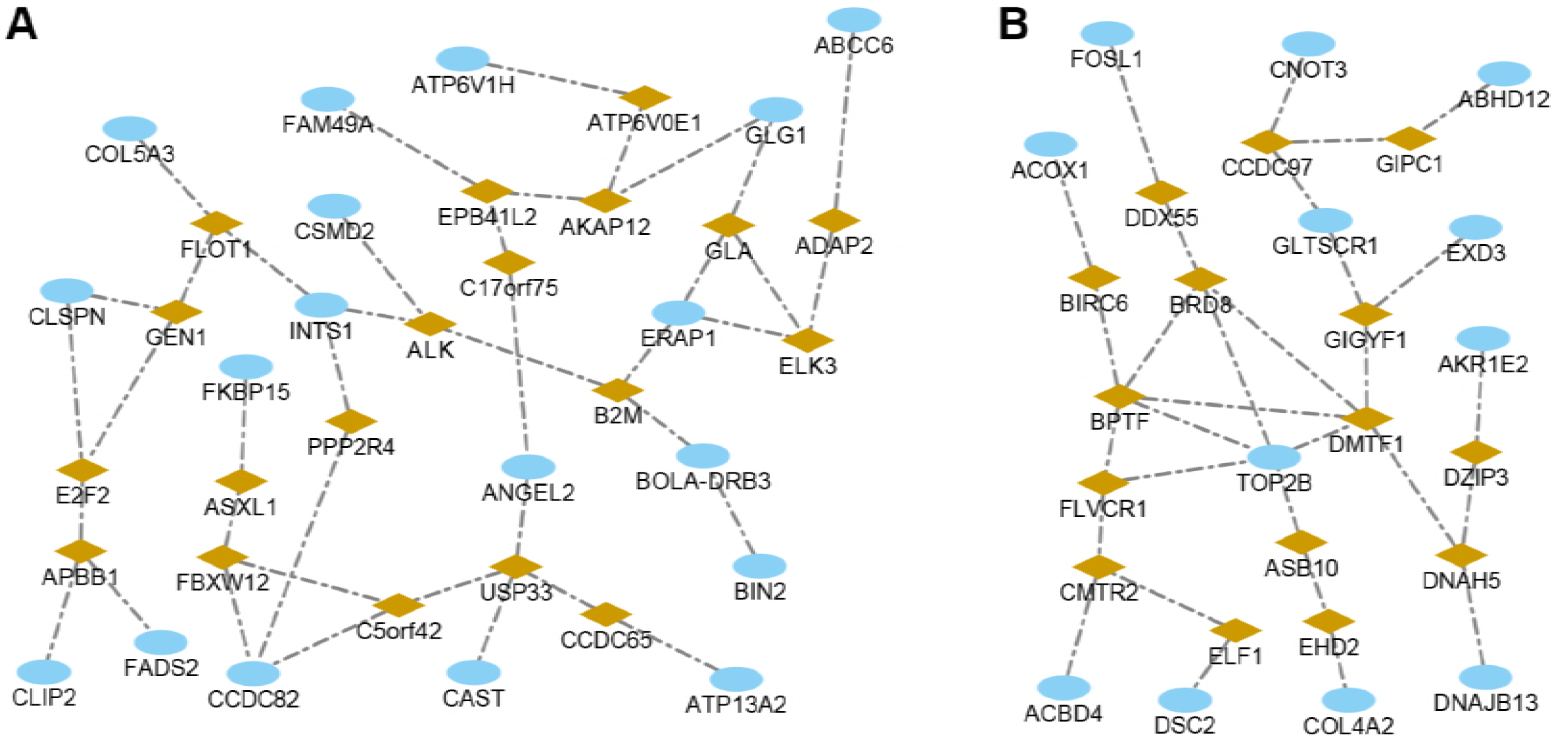
The respective sub-networks for the genes shared by the higher phenotype groups and the lower phenotype groups of body length and body girth in cattle. (A) the sub-network extracted from 62 genes shared by higher phenotype value in body girth group (BG_H) and body length (BL_H); (B) the sub-network extracted from 66 genes shared by lower phenotype value in body girth (BG_L) and body length (BL_L). The blue circles were the genes with rare variants in our data. The orange diamond shapes were the linker genes to connect those mutated genes for a fully-connected network.

## 4. Discussion

Growth traits like body height, length and girth are classic quantitative traits, reflecting the combined influence of multiple polygenic factors. Thus, the study of growth traits is an ideal opportunity to dissect the architecture of a highly polygenic trait in human, animal or livestock. Several common variants associated with phenotypic variation in human height were detected, such as two variants in the region of *HMGA2* (Weedon et al., 2007) and *GDF5-UQCC* (Sanna et al., 2008). Moreover, a meta-GWAS analysis for human height identified ten newly loci strongly associated with variation in height, and found several pathways, like let-7 targets and Hedgehog signalling, as important regulators of human stature (Lettre et al., 2008). However, these significant loci together only account for around 2% of the population variation in height together. Similarly another study found that hundreds of variants clustered in genomic loci also explained no more than 10% phenotypic variation (Lango Allen et al., 2010). On the other hand, the same researches on dissection the genetic variants of body size traits have been conducted on livestock like sheep (Berenos et al., 2015), cattle (Gutierrez-Gil et al., 2009), pig (Guo et al., 2015). Numerous candidate regions and genes have been screened. For instance, the candidate gene *NCAPG* where a non-synonymous but chemically conserved variant was proposed to be a potential causative variant for body frame size in cattle (Setoguchi et al., 2011). Similarly, SNPs resided in *NCAPG* and *LCORL* genes have also been reported to be associated with several body size traits for horse breeding (Signer-Hasler et al., 2012). Overall, GWAS studies have identified large numbers of loci and variants that implicate biologically relevant genes and pathways of growth traits; however, they mainly focused on the common genetic variants and explained a limited genetic contribution. Thus, additional approaches are needed to fully dissect the genetic architecture of growth traits. Traditionally, growth traits are included in selection criteria of beef cattle breeding programs, and specifically body length and girth have been shown to be a useful predictors for cattle liveweight (Enevoldsen and Kristensen, 1997) (Lukuyu et al., 2016). Considering that our GWAS results for five growth traits (body length, height, hip height, heart girth, and abdominal girth) have no significant loci detected (*P* < 5×10^−7^) in beef cattle, we used exome sequencing analysis to capture the rare and novel variants associated with body length and girth.

Through the gene annotation and functional enrichment analysis for candidate rare variants in four trait groups (BG_H, BG_L, BL_H and BL_L), 886 unique genes in four trait groups were found to be associated with various developmental processes and organelle organization related functional terms. Interestingly, multiple genetic loci for body size were also confirmed to be associated with developmental pathways by using GWAS studies in Chinese Holstein cattle (Zhang et al., 2017). We also identified a number of organelle organization related functional terms. For Eukaryotic organisms, there is precise regulatory mechanism on their growth across the diverse length scales of biological organization. Recent studies in intestine of *C. elegans* worm revealed that the volume of the nucleolus is directly proportional (isometric) to cell size during larval development (Uppaluri et al., 2016). Furthermore, the relative size of the nucleolus is predictive of the growth rate of the entire worm. Similarly, our result suggested that the growth in organelle level could be involved in the growth in tissue and body development.

In addition, six significantly over-represented pathways were identified by KEGG pathway analysis for these genes, which are mainly related to the Collagen formation and biosynthesis, Collagen and modifying enzymes, and Collagen fibrils assembly. Previously studies reported that the diameter of collagen fibrils could increase gradually during embryogenesis, tissue growth and body size (Pilotto and Filosi, 1977). Recent molecular mechanism studies revealed that hydrolysed collagen intake could increase bone mass of growing in rats (Takeda et al., 2013). Combined with the results from GO enrichment analysis, our results highlight the potential importance of collagen formation and organelle organization for body size growth in multicellular organisms.

Since body girth has highly positive genetic correlation with body length, the analysis of those genetic variants shared by same level (higher or lower) of BG and BL samples may help to further identify candidate genes associated with growth and gain of weight in cattle. As shown in Figure 5, we reconstructed sub-network specific for 62, 66 genes shared by BL_H and BG_H, BL_L and BG_L group respectively to investigate their gene co-expressions. In the sub-network of high phenotype in BL_H and BG_H groups, the *INTS1*, a subunit of the integrator complex and mediated 3-prime end processing of small nuclear RNAs U1 and U2, was co-expressed with *ALK*, which plays an important role in the cell growth and brain development, and exerts its effects on specific neurons in the nervous system.(Motegi et al., 2004) We also found a linker gene *B2M* (Beta 2-Microglobulin), a component of the major histocompatibility complex in chordates, which has growth factor-like activity for cultured rat cells (Centrella et al., 1989) and thus may have an impact on the body length and girth of cattle. Meanwhile, another calpastatin gene *CAST* mapped to BTA7 was also linked to multiple growth-related traits in view of our QTL-based analysis (Table S6_a). Basically, previous studies suggested that *CAST* was associated with insulin-like growth factor 1 (*IGF-1*) levels and body weight, which may imply its key role in the body growth (Pintos and Corva, 2011). Another GWAS study proposed *CAST* gene also as a functional and positional candidate gene for carcass and meat quality traits in beef cattle (Curi et al., 2010; Curi et al., 2009). In general, comparing to GWAS-based association studies, our rare variants-based approach is powerful to identify genes directly affect phenotype at the molecular level, rather than merely conferring the association or risk. Therefore, these genes may have potential direct links to the body size control at the cellular level. Moreover, we focused on the network for the 66 genes shared by the lower phenotype in BL_L and BG_L groups, and found two genes (*DSC2* and *DZIP3*) associated with multiple reproductive traits such as conception rate and daughter pregnancy rate (Table S6_b). Additionally, the putative candidate gene *TOP2B* is known to be associated with various neurodevelopmental conditions (Harkin et al., 2016), and a recent study suggested that *de novo TOP2B* mutation may lead to global developmental delay and intellectual disability (Lam et al., 2017). In summary, our study has identified many co-expressed genes with genetic rare mutations. A number of these genes are turned out to be associated with other complex traits, which suggest their potential links to phenotypes.

In conclusion, we have presented the first whole exome-capture sequencing study of body girth and length for Simmental cattle and the network view of genetic changes against the whole proteome in cattle. Our study provides not only a comprehensive genetic resource of beef cattle for the effective breeding but also illustrated a comprehensive mutational catalogue for body growth-related trait. The identified genes may reveal the importance of collagen formation and organelle organization for body size growth in multicellular organisms. By analysis the genes with rare nonsynonymous variants, we identified several genes with association to body growth and reproduction.

## Availability

The sequence data reported in this study have been deposited in the genome sequence archive of Beijing Institute of Genomics, Chinese Academy of Sciences (gsa.big.ac.cn) under the accession no. PRJCA000519, PRJCA000515, PRJCA000513, and PRJCA000510.

## Acknowledgments

This work was supported by grants from the National Natural Science Foundation of China (31402039), Cattle Breeding Innovative Research Team of Chinese Academy of Agricultural Sciences (cxgc-ias-03), the National Beef Cattle Industrial Technology System (CARS-37), China Scholarship Council (CSC), and the start-up grant to Dr. Min Zhao.

## Supplementary data

**Supplementary Table S1. Sample information.**

**Supplementary Table S2. All the genetic variants shared by all the samples in the same trait groups.** (a) BG_H, (b) BG_L, (c) BL_H, (d) BL_L.

**Supplementary Table S3. The 886 unique genes associated with rare nonsynonymous variants from all the samples.**

**Supplementary Table S4. The GO (gene ontology) and KEGG pathway enrichment analysis of 886 genes with rare nonsynonymous variants.**

**Supplementary Table S5. Rare variants shared by different phenotypic groups.** (a) The 69 rare variants shared by two higher phenotype groups in body girth (BG_H) and body length (BL_H). (b) The 67 rare variants shared by two lower phenotype groups in body girth (BG_L) and body length (BL_L).

**Supplementary Table S6. Cattle QTL information from animal QTL database.** (a) The QTL analysis of 62 genes associated with higher phenotype in body girth (BG_H) and body length (BL_H). (b) The QTL analysis of 66 genes associated with lower phenotype in body girth (BG_L) and body length (BL_L).

## References

Beiki, H., Nejati-Javaremi, A., Pakdel, A., Masoudi-Nejad, A., Hu, Z.L., Reecy, J.M., 2016. Large-scale gene co-expression network as a source of functional annotation for cattle genes. Bmc Genomics 17.

Berenos, C., Ellis, P.A., Pilkington, J.G., Lee, S.H., Gratten, J., Pemberton, J.M., 2015. Heterogeneity of genetic architecture of body size traits in a free-living population. Mol Ecol 24, 1810–1830.

Bush, W.S., Moore, J.H., 2012. Chapter 11: Genome-Wide Association Studies. Plos Comput Biol 8.

Centrella, M., McCarthy, T., Canalis, E., 1989. Beta 2-microglobulin enhances insulin-like growth factor I receptor levels and synthesis in bone cell cultures. Journal of Biological Chemistry 264, 18268–18271.

Chen, Y., Liu, Y.N., Du, M., Zhang, W.G., Xu, L., Gao, X., Zhang, L.P., Gao, H.J., Xu, L.Y., Li, J.Y., Zhao, M., 2017. Constructing a comprehensive gene co-expression based interactome in Bos taurus. Peerj 5.

Clayton, D.G., Walker, N.M., Smyth, D.J., Pask, R., Cooper, J.D., Maier, L.M., Smink, L.J., Lam, A.C., Ovington, N.R., Stevens, H.E., Nutland, S., Howson, J.M.M., Faham, M., Moorhead, M., Jones, H.B., Falkowski, M., Hardenbol, P., Willis, T.D., Todd, J.A., 2005. Population structure, differential bias and genomic control in a large-scale, case-control association study. Nat Genet 37, 1243–1246.

Curi, R.A., Chardulo, L.A., Giusti, J., Silveira, A.C., Martins, C.L., de Oliveira, H.N., 2010. Assessment of GH1, CAPN1 and CAST polymorphisms as markers of carcass and meat traits in Bos indicus and Bos taurus-Bos indicus cross beef cattle. Meat Sci 86, 915–920.

Curi, R.A., Chardulo, L.A., Mason, M.C., Arrigoni, M.D., Silveira, A.C., de Oliveira, H.N., 2009. Effect of single nucleotide polymorphisms of CAPN1 and CAST genes on meat traits in Nellore beef cattle (Bos indicus) and in their crosses with Bos taurus. Anim Genet 40, 456–462.

Donnelly, P., 2008. Progress and challenges in genome-wide association studies in humans. Nature 456, 728–731.

Elsik, C.G., Unni, D.R., Diesh, C.M., Tayal, A., Emery, M.L., Nguyen, H.N., Hagen, D.E., 2016. Bovine Genome Database: new tools for gleaning function from the Bos taurus genome. Nucleic Acids Res 44, D834–839.

Enevoldsen, C., Kristensen, T., 1997. Estimation of body weight from body size measurements and body condition scores in dairy cows. J Dairy Sci 80, 1988–1995.

Fan, H., Wu, Y., Zhou, X., Xia, J., Zhang, W., Song, Y., Liu, F., Chen, Y., Zhang, L., Gao, X., Gao, H., Li, J., 2015. Pathway-Based Genome-Wide Association Studies for Two Meat Production Traits in Simmental Cattle. Sci Rep 5, 18389.

Ghorbani, S., Tahmoorespur, M., Nejad, A.M., Nasiri, M.R., Asgari, Y., 2015. Analysis of the enzyme network involved in cattle milk production using graph theory. Mol Biol Res Commun 4, 93–103.

Guo, Y., Hou, L., Zhang, X., Huang, M., Mao, H., Chen, H., Ma, J., Chen, C., Ai, H., Ren, J., Huang, L., 2015. A meta analysis of genome-wide association studies for limb bone lengths in four pig populations. BMC Genet 16, 95.

Gutierrez-Gil, B., Williams, J.L., Homer, D., Burton, D., Haley, C.S., Wiener, P., 2009. Search for quantitative trait loci affecting growth and carcass traits in a cross population of beef and dairy cattle. J Anim Sci 87, 24–36.

Harkin, L.F., Gerrelli, D., Gold Diaz, D.C., Santos, C., Alzu’bi, A., Austin, C.A., Clowry, G.J., 2016. Distinct expression patterns for type II topoisomerases IIA and IIB in the early foetal human telencephalon. Journal of anatomy 228, 452–463.

Hirschhorn, J.N., Daly, M.J., 2005. Genome-wide association studies for common diseases and complex traits. Nat Rev Genet 6, 95–108.

Hu, Z.L., Park, C.A., Wu, X.L., Reecy, J.M., 2013. Animal QTLdb: an improved database tool for livestock animal QTL/association data dissemination in the post-genome era. Nucleic Acids Res 41, D871–879.

Jahuey-Martinez, F.J., Parra-Bracamonte, G.M., Sifuentes-Rincon, A.M., Martinez-Gonzalez, J.C., Gondro, C., Garcia-Perez, C.A., Lopez-Bustamante, L.A., 2016. Genomewide association analysis of growth traits in Charolais beef cattle. J Anim Sci 94, 4570–4582.

Jiang, Y., Ying, W.T., Wu, S.F., Chen, M., Guan, W., Yang, D., Song, Y.P., Liu, X., Li, J.Q., Hao, Y.W., Sun, A.H., Geng, C., Li, H., Mi, W., Zhang, Y.J., Zhang, J.Y., Chen, X.L., Li, L., Gong, Y., Li, T., Ma, J., Li, D., Yuan, X.Y., Zhang, X.Q., Xue, X.F., Zhu, Y.P., Qian, X.H., He, F.C., Zhong, F., Shen, H.L., Lin, C.Z., Lu, H.J., Liu, X.H., Wei, L.M., Cao, J., Yun, D., Zhang, J., Gao, M.X., Fan, H.Z., Zhang, Y., Cheng, G., Yu, Y.Y., Xie, L.Q., Wang, H., Zhang, X.M., He, F.C., Yang, P.Y., Shi, L., Tong, W., Li, X.L., Wang, Y., Liu, S.Q., Sheng, Q.H., Zeng, R., Sun, Y.L., Xu, Y., Cai, J.Q., He, P., Gao, H.J., Zhao, X.H., Tan, Y.X., Yan, H.X., Yang, Y., Wang, H.Y., Huang, J., Han, Z.G., He, Q.Y., Chen, P., Liang, S.P., Zhao, M., Mao, X.Z., Wei, L.P., Yu, H., Cao, Z.W., Li, Y.X., Dai, W.J., Jiang, H.C., Wang, D.G., Zheng, J.J., Gao, X., Tang, Y., Li, X.Y., Cheng, J.P., Liu, Y.H., Zhang, X.N., Wang, X.F., Jia, J.D., An, D.C., Wang, Z., Li, Q., Cui, T., Proteome, C.H.L., 2010. First Insight into the Human Liver Proteome from PROTEOMESKY-LIVERHu 1.0, a Publicly Available Database. J Proteome Res 9, 79–94.

Kong, L., Cheng, L., Fan, L.Y., Zhao, M., Qu, H., 2013. IQdb: an intelligence quotient score-associated gene resource for human intelligence. Database (Oxford) 2013, bat063.

Lam, C.W., Yeung, W.L., Law, C.Y., 2017. Global developmental delay and intellectual disability associated with a de novo TOP2B mutation. Clin Chim Acta 469, 63–68.

Lango Allen, H., Estrada, K., Lettre, G., Berndt, S.I., Weedon, M.N., Rivadeneira, F., Willer, C.J., Jackson, A.U., Vedantam, S., Raychaudhuri, S., Ferreira, T., Wood, A.R., Weyant, R.J., Segre, A.V., Speliotes, E.K., Wheeler, E., Soranzo, N., Park, J.H., Yang, J., Gudbjartsson, D., Heard-Costa, N.L., Randall, J.C., Qi, L., Vernon Smith, A., Magi, R., Pastinen, T., Liang, L., Heid, I.M., Luan, J., Thorleifsson, G., Winkler, T.W., Goddard, M.E., Sin Lo, K., Palmer, C., Workalemahu, T., Aulchenko, Y.S., Johansson, A., Zillikens, M.C., Feitosa, M.F., Esko, T., Johnson, T., Ketkar, S., Kraft, P., Mangino, M., Prokopenko, I., Absher, D., Albrecht, E., Ernst, F., Glazer, N.L., Hayward, C., Hottenga, J.J., Jacobs, K.B., Knowles, J.W., Kutalik, Z., Monda, K.L., Polasek, O., Preuss, M., Rayner, N.W., Robertson, N.R., Steinthorsdottir, V., Tyrer, J.P., Voight, B.F., Wiklund, F., Xu, J., Zhao, J.H., Nyholt, D.R., Pellikka, N., Perola, M., Perry, J.R., Surakka, I., Tammesoo, M.L., Altmaier, E.L., Amin, N., Aspelund, T., Bhangale, T., Boucher, G., Chasman, D.I., Chen, C., Coin, L., Cooper, M.N., Dixon, A.L., Gibson, Q., Grundberg, E., Hao, K., Juhani Junttila, M., Kaplan, L.M., Kettunen, J., Konig, I.R., Kwan, T., Lawrence, R.W., Levinson, D.F., Lorentzon, M., McKnight, B., Morris, A.P., Muller, M., Suh Ngwa, J., Purcell, S., Rafelt, S., Salem, R.M., Salvi, E., Sanna, S., Shi, J., Sovio, U., Thompson, J.R., Turchin, M.C., Vandenput, L., Verlaan, D.J., Vitart, V., White, C.C., Ziegler, A., Almgren, P., Balmforth, A.J., Campbell, H., Citterio, L., De Grandi, A., Dominiczak, A., Duan, J., Elliott, P., Elosua, R., Eriksson, J.G., Freimer, N.B., Geus, E.J., Glorioso, N., Haiqing, S., Hartikainen, A.L., Havulinna, A.S., Hicks, A.A., Hui, J., Igl, W., Illig, T., Jula, A., Kajantie, E., Kilpelainen, T.O., Koiranen, M., Kolcic, I., Koskinen, S., Kovacs, P., Laitinen, J., Liu, J., Lokki, M.L., Marusic, A., Maschio, A., Meitinger, T., Mulas, A., Pare, G., Parker, A.N., Peden, J.F., Petersmann, A., Pichler, I., Pietilainen, K.H., Pouta, A., Ridderstrale, M., Rotter, J.I., Sambrook, J.G., Sanders, A.R., Schmidt, C.O., Sinisalo, J., Smit, J.H., Stringham, H.M., Bragi Walters, G., Widen, E., Wild, S.H., Willemsen, G., Zagato, L., Zgaga, L., Zitting, P., Alavere, H., Farrall, M., McArdle, W.L., Nelis, M., Peters, M.J., Ripatti, S., van Meurs, J.B., Aben, K.K., Ardlie, K.G., Beckmann, J.S., Beilby, J.P., Bergman, R.N., Bergmann, S., Collins, F.S., Cusi, D., den Heijer, M., Eiriksdottir, G., Gejman, P.V., Hall, A.S., Hamsten, A., Huikuri, H.V., Iribarren, C., Kahonen, M., Kaprio, J., Kathiresan, S., Kiemeney, L., Kocher, T., Launer, L.J., Lehtimaki, T., Melander, O., Mosley, T.H., Jr., Musk, A.W., Nieminen, M.S., O’Donnell, C.J., Ohlsson, C., Oostra, B., Palmer, L.J., Raitakari, O., Ridker, P.M., Rioux, J.D., Rissanen, A., Rivolta, C., Schunkert, H., Shuldiner, A.R., Siscovick, D.S., Stumvoll, M., Tonjes, A., Tuomilehto, J., van Ommen, G.J., Viikari, J., Heath, A.C., Martin, N.G., Montgomery, G.W., Province, M.A., Kayser, M., Arnold, A.M., Atwood, L.D., Boerwinkle, E., Chanock, S.J., Deloukas, P., Gieger, C., Gronberg, H., Hall, P., Hattersley, A.T., Hengstenberg, C., Hoffman, W., Lathrop, G.M., Salomaa, V., Schreiber, S., Uda, M., Waterworth, D., Wright, A.F., Assimes, T.L., Barroso, I., Hofman, A., Mohlke, K.L., Boomsma, D.I., Caulfield, M.J., Cupples, L.A., Erdmann, J., Fox, C.S., Gudnason, V., Gyllensten, U., Harris, T.B., Hayes, R.B., Jarvelin, M.R., Mooser, V., Munroe, P.B., Ouwehand, W.H., Penninx, B.W., Pramstaller, P.P., Quertermous, T., Rudan, I., Samani, N.J., Spector, T.D., Volzke, H., Watkins, H., Wilson, J.F., Groop, L.C., Haritunians, T., Hu, F.B., Kaplan, R.C., Metspalu, A., North, K.E., Schlessinger, D., Wareham, N.J., Hunter, D.J., O’Connell, J.R., Strachan, D.P., Wichmann, H.E., Borecki, I.B., van Duijn, C.M., Schadt, E.E., Thorsteinsdottir, U., Peltonen, L., Uitterlinden, A.G., Visscher, P.M., Chatterjee, N., Loos, R.J., Boehnke, M., McCarthy, M.I., Ingelsson, E., Lindgren, C.M., Abecasis, G.R., Stefansson, K., Frayling, T.M., Hirschhorn, J.N., 2010. Hundreds of variants clustered in genomic loci and biological pathways affect human height. Nature 467, 832–838.

Lettre, G., Jackson, A.U., Gieger, C., Schumacher, F.R., Berndt, S.I., Sanna, S., Eyheramendy, S., Voight, B.F., Butler, J.L., Guiducci, C., Illig, T., Hackett, R., Heid, I.M., Jacobs, K.B., Lyssenko, V., Uda, M., Diabetes Genetics, I., Fusion, Kora, Prostate, L.C., Ovarian Cancer Screening, T., Nurses’ Health, S., SardiNia, Boehnke, M., Chanock, S.J., Groop, L.C., Hu, F.B., Isomaa, B., Kraft, P., Peltonen, L., Salomaa, V., Schlessinger, D., Hunter, D.J., Hayes, R.B., Abecasis, G.R., Wichmann, H.E., Mohlke, K.L., Hirschhorn, J.N., 2008. Identification of ten loci associated with height highlights new biological pathways in human growth. Nat Genet 40, 584–591.

Li, H., Handsaker, B., Wysoker, A., Fennell, T., Ruan, J., Homer, N., Marth, G., Abecasis, G., Durbin, R., 2009. The Sequence Alignment/Map format and SAMtools. Bioinformatics 25, 2078–2079.

Lukuyu, M.N., Gibson, J.P., Savage, D.B., Duncan, A.J., Mujibi, F.D.N., Okeyo, A.M., 2016. Use of body linear measurements to estimate liveweight of crossbred dairy cattle in smallholder farms in Kenya. Springerplus 5.

Motegi, A., Fujimoto, J., Kotani, M., Sakuraba, H., Yamamoto, T., 2004. ALK receptor tyrosine kinase promotes cell growth and neurite outgrowth. J Cell Sci 117, 3319–3329.

Pertille, F., Moreira, G.C.M., Zanella, R., Nunes, J.D.D., Boschiero, C., Rovadoscki, G.A., Mourao, G.B., Ledur, M.C., Coutinho, L.L., 2017. Genome-wide association study for performance traits in chickens using genotype by sequencing approach. Sci Rep 7, 11.

Pilotto, F., Filosi, M., 1977. Relationship between collagen fibril diameters and body size. Study of fish derma. Cell Tissue Res 182, 119–131.

Pintos, D., Corva, P.M., 2011. Association between molecular markers for beef tenderness and growth traits in Argentinian angus cattle. Anim Genet 42, 329–332.

Sanna, S., Jackson, A.U., Nagaraja, R., Willer, C.J., Chen, W.M., Bonnycastle, L.L., Shen, H., Timpson, N., Lettre, G., Usala, G., Chines, P.S., Stringham, H.M., Scott, L.J., Dei, M., Lai, S., Albai, G., Crisponi, L., Naitza, S., Doheny, K.F., Pugh, E.W., Ben-Shlomo, Y., Ebrahim, S., Lawlor, D.A., Bergman, R.N., Watanabe, R.M., Uda, M., Tuomilehto, J., Coresh, J., Hirschhorn, J.N., Shuldiner, A.R., Schlessinger, D., Collins, F.S., Davey Smith, G., Boerwinkle, E., Cao, A., Boehnke, M., Abecasis, G.R., Mohlke, K.L., 2008. Common variants in the GDF5-UQCC region are associated with variation in human height. Nat Genet 40, 198–203.

Setoguchi, K., Watanabe, T., Weikard, R., Albrecht, E., Kuhn, C., Kinoshita, A., Sugimoto, Y., Takasuga, A., 2011. The SNP c.1326T>G in the non-SMC condensin I complex, subunit G (NCAPG) gene encoding a p.Ile442Met variant is associated with an increase in body frame size at puberty in cattle. Anim Genet 42, 650–655.

Shannon, P., Markiel, A., Ozier, O., Baliga, N.S., Wang, J.T., Ramage, D., Amin, N., Schwikowski, B., Ideker, T., 2003. Cytoscape: A software environment for integrated models of biomolecular interaction networks. Genome Research 13, 2498–2504.

Signer-Hasler, H., Flury, C., Haase, B., Burger, D., Simianer, H., Leeb, T., Rieder, S., 2012. A genome-wide association study reveals loci influencing height and other conformation traits in horses. PLoS One 7, e37282.

Sorbolini, S., Bongiorni, S., Cellesi, M., Gaspa, G., Dimauro, C., Valentini, A., Macciotta, N.P.P., 2017. Genome wide association study on beef production traits in Marchigiana cattle breed. J Anim Breed Genet 134, 43–48.

Takeda, S., Park, J.H., Kawashima, E., Ezawa, I., Omi, N., 2013. Hydrolyzed collagen intake increases bone mass of growing rats trained with running exercise. J Int Soc Sports Nutr 10, 35.

Uppaluri, S., Weber, S.C., Brangwynne, C.P., 2016. Hierarchical Size Scaling during Multicellular Growth and Development. Cell Rep 17, 345–352.

van den Berg, I., Boichard, D., Guldbrandtsen, B., Lund, M.S., 2016. Using Sequence Variants in Linkage Disequilibrium with Causative Mutations to Improve Across-Breed Prediction in Dairy Cattle: A Simulation Study. G3 (Bethesda) 6, 2553–2561.

Weedon, M.N., Lettre, G., Freathy, R.M., Lindgren, C.M., Voight, B.F., Perry, J.R., Elliott, K.S., Hackett, R., Guiducci, C., Shields, B., Zeggini, E., Lango, H., Lyssenko, V., Timpson, N.J., Burtt, N.P., Rayner, N.W., Saxena, R., Ardlie, K., Tobias, J.H., Ness, A.R., Ring, S.M., Palmer, C.N., Morris, A.D., Peltonen, L., Salomaa, V., Diabetes Genetics, I., Wellcome Trust Case Control, C., Davey Smith, G., Groop, L.C., Hattersley, A.T., McCarthy, M.I., Hirschhorn, J.N., Frayling, T.M., 2007. A common variant of HMGA2 is associated with adult and childhood height in the general population. Nat Genet 39, 1245–1250.

Xia, J., Qi, X., Wu, Y., Zhu, B., Xu, L., Zhang, L., Gao, X., Chen, Y., Li, J., Gao, H., 2016. Genome-wide association study identifies loci and candidate genes for meat quality traits in Simmental beef cattle. Mamm Genome 27, 246–255.

Xia, J.W., Fan, H.Z., Chang, T.P., Xu, L.Y., Zhang, W.G., Song, Y.X., Zhu, B., Zhang, L.P., Gao, X., Chen, Y., Li, J.Y., Gao, H.J., 2017. Searching for new loci and candidate genes for economically important traits through genebased association analysis of Simmental cattle. Sci Rep 7, 9.

Zhang, Q., Guldbrandtsen, B., Calus, M.P., Lund, M.S., Sahana, G., 2016a. Comparison of gene-based rare variant association mapping methods for quantitative traits in a bovine population with complex familial relationships. Genet Sel Evol 48, 60.

Zhang, W., Li, J., Guo, Y., Zhang, L., Xu, L., Gao, X., Zhu, B., Gao, H., Ni, H., Chen, Y., 2016b. Multi-strategy genome-wide association studies identify the DCAF16-NCAPG region as a susceptibility locus for average daily gain in cattle. Sci Rep 6, 38073.

Zhang, W., Xu, L., Gao, H., Wu, Y., Gao, X., Zhang, L., Zhu, B., Song, Y., Bao, J., Li, J., Chen, Y., 2016c. Detection of candidate genes for growth and carcass traits using genome-wide association strategy in Chinese Simmental beef cattle. Animal Production Science, -.

Zhang, X., Chu, Q., Guo, G., Dong, G., Li, X., Zhang, Q., Zhang, S., Zhang, Z., Wang, Y., 2017. Genome-wide association studies identified multiple genetic loci for body size at four growth stages in Chinese Holstein cattle. PLoS One 12, e0175971.

Zhao, M., Chen, X., Gao, G., Tao, L., Wei, L., 2009. RLEdb: a database of rate-limiting enzymes and their regulation in human, rat, mouse, yeast and E. coli. Cell Res 19, 793–795.

Zhao, M., Li, X., Qu, H., 2013. EDdb: a web resource for eating disorder and its application to identify an extended adipocytokine signaling pathway related to eating disorder. Sci China Life Sci 56, 1086–1096.

Zimin, A.V., Delcher, A.L., Florea, L., Kelley, D.R., Schatz, M.C., Puiu, D., Hanrahan, F., Pertea, G., Van Tassell, C.P., Sonstegard, T.S., Marcais, G., Roberts, M., Subramanian, P., Yorke, J.A., Salzberg, S.L., 2009. A whole-genome assembly of the domestic cow, Bos taurus. Genome Biol 10, R42.

